# Detecting Sleep Deprivation from Running Biomechanics Using Machine Learning Classification: A Comparison Between Wearable and Laboratory Motion Capture

**DOI:** 10.64898/2026.07.14.738397

**Authors:** Maura Seynaeve, Kilian Hendrickx, Benedicte Vanwanseele, Toon de Beukelaar

## Abstract

Sleep deprivation is associated with impaired endurance performance and an increased risk of running-related injury. Previous research has identified alterations in running biomechanics following a single night of sleep deprivation under laboratory conditions. However, whether these biomechanical changes can be detected using wearable technology remains unknown. Twenty-one recreationally active runners completed submaximal treadmill running under both normal sleep and total sleep deprivation conditions in a randomized crossover design. Biomechanical features were extracted simultaneously using a full-body motion capture system and a trunk-mounted wearable sensor. Five machine learning classifiers were evaluated in two classification tasks: a within-subject task using paired recordings from the same individual, and a between-subject task performed without individual baseline data. Within-subject classification consistently exceeded chance level for both measurement systems, with best accuracies of 85% for the wearable sensor (Logistic Regression) and 83% for the motion capture system (Random Forest). These findings indicate that sleep deprivation produces a systematic and individually consistent biomechanical signature during running. In contrast, between-subject classification failed across nearly all models and systems, with accuracies remaining close to chance level (∼50%), demonstrating that inter-individual variability obscures the sleep-deprivation signal in the absence of personalized baseline data. Both systems converged on temporal organization, loading-related variables, and stride-to-stride variability as the most discriminative feature domains. Contrary to expectations, the laboratory motion capture system did not outperform the wearable sensor. Together, these findings demonstrate that individualized, baseline-referenced monitoring is essential for detecting sleep-deprivation-related changes in running gait, and suggest that a single trunk-mounted wearable sensor may provide a practical solution for real-world monitoring when paired recordings are available.

## 1 Introduction

Recreational running is one of the most popular forms of physical activity worldwide, with millions of people participating regularly across all ages and fitness levels (Hulteen et al., 2017). While running offers well-established health benefits (Lee et al., 2014), it carries a substantial injury burden. Previous research suggests that as many as 65% of runners experience at least one overuse injury each year, with common examples including patellofemoral pain syndrome, iliotibial band friction syndrome, and medial tibial stress syndrome(Dias Lopes et al., 2012; Messier et al., 2018). These injuries arise from the repetitive mechanical loading inherent to running, where cumulative stress applied to muscles, tendons, and joints may eventually exceed a structure’s tolerance threshold(Bertelsen et al., 2017). Importantly, fatigue is a key modulator of this process: as fatigue develops, running biomechanics change in ways that can introduce atypical loading patterns, potentially pushing tissues beyond their limits (Zandbergen et al., 2023). Because these biomechanical alterations are often subtle, objective monitoring tools are essential for coaches and clinicians working with this population (Mason et al., 2022).

When assessing exercise-induced fatigue, laboratory-based techniques have frequently been used; however, wearable sensors have more recently emerged as a practical tool for detecting such fatigue, defined as the acute, task-related decline in performance during or following a sustained effort (Marotta et al., 2022). Evidence from these studies shows that, in running, peak tibial and sacral acceleration increases with fatigue, suggesting greater impact loading as fatigue develops (Schütte et al., 2015; Reenalda et al., 2016, 2019). Changes in shock attenuation between body segments have also been reported, with several studies observing increased head-to-tibia and sacrum-to-tibia attenuation under fatigued conditions, indicating altered impact transmission strategies(Derrick et al., 2002; Encarnación-Martínez et al., 2023). Beyond impact-related variables, only a limited number of studies have reported fatigue-related changes in spatiotemporal gait parameters such as stride length, stride frequency, and contact time, though these findings are less consistent (Schütte et al., 2015; Reenalda et al., 2016; Meyer et al., 2021). Collectively, this body of work demonstrates that wearable accelerometers are sensitive to fatigue-induced biomechanical shifts during running.

However, fatigue is a multifactorial phenomenon that can arise from a variety of physiological and psychological stressors. Alongside exercise-induced fatigue, which arises from the accumulation of physiological and neuromuscular stress during prolonged or intense exercise, insufficient sleep represents a clinically important and underexplored source of fatigue for running populations. Insufficient sleep has been associated with impaired athletic performance and injury-risk in athletes, making sleep health an important concern for coaches and clinicians working with this population (Charest and Grandner, 2022). A recent meta-analysis of 31 studies showed that sleep deprivation has a moderate negative effect on endurance performance, with longer-duration exercise (>30 min) being more adversely affected (Lopes et al., 2023). In addition, a 26-week prospective cohort study of 339 runners found that poorer sleep quality was associated with a 36% higher risk of running-related injury (Goldberg et al., 2025). Despite these effects on running-related injury, studies examining the direct biomechanical consequences of fatigue induced by sleep deprivation remain scarce.

The biomechanical consequences of fatigue induced by sleep loss cannot be assumed equivalent to the consequences of exercise-induced fatigue as described above. Unlike the peripheral muscular fatigue caused by metabolite accumulation or energy substrate depletion, sleep deprivation is more likely to influence motor performance through central mechanisms, occurring proximal to the neuromuscular junction. Beyond its well-documented effects on cognitive performance(Killgore, 2010), sleep loss has been shown to impair motor performance in a task-dependent manner (Fullagar et al., 2015; Knowles et al., 2018; Vitale et al., 2019; Walsh et al., 2021; Craven et al., 2022). In general, effects are more pronounced for continuous, multi-joint movements than for discrete, single-joint tasks (Fullagar et al., 2015; Knowles et al., 2018; Vitale et al., 2019; Walsh et al., 2021; Craven et al., 2022). This distinction is mechanistically meaningful: complex, continuous movements such as maintaining balance or walking rely on higher-order neural control processes, including feedforward planning, sensorimotor feedback, and executive attention, all of which are sensitive to sleep loss (Woollacott and Shumway-Cook, 2002; Pruszynski et al., 2011; Stuhr et al., 2018; Umemura et al., 2022). Running, as a dynamic, rhythmically coordinated, multi-joint task, is therefore likely to be vulnerable to these centrally mediated disruptions. Supporting this, recent work using a marker-based 3D motion capture system and instrumented treadmill confirmed meaningful biomechanical changes following a single night of sleep deprivation (Seynaeve et al., 2026). However, such laboratory-grade equipment is rarely accessible outside of research settings. Hence, this raises the question whether these alterations could instead be detected using wearable technology, a more scalable and ecologically valid alternative.

A frequent challenge in detecting these condition-induced biomechanical patterns is that inter-individual differences in gait patterns routinely exceed the magnitude of condition effects, causing group-level analyses to wash out the very signal of interest. As has been demonstrated in the exercise-induced fatigue literature, the average runner does not resemble any individual in the group, and population-level models frequently mask meaningful individual responses (Chalitsios et al., 2024). Machine learning has emerged as a promising framework for addressing this problem, precisely because it captures coordinated patterns across many biomechanical variables simultaneously rather than testing each in isolation. Early research established the feasibility of this approach for exercise-induced fatigue detection. For instance, Buckley et al. demonstrated that a single inertial measurement unit (IMU) placed at the lumbar spine could classify non-fatigued versus fatigued states using a single group-level model with an accuracy of 75% (Buckley et al., 2017). Additional work showed that this could be improved substantially through individualization. Subject-specific models consistently and significantly outperform group-based equivalents in identifying biomechanical shifts (Buckley et al., 2017; Op De Beéck et al., 2018; Dimmick et al., 2023). This performance gap exists because the most discriminative gait variables are highly subject-dependent; for instance, the specific mechanical features that signal fatigue or environmental adaptation in one runner often hold little predictive power for another (Marotta et al., 2021). To reliably isolate these subtle shifts, successful models typically require normalizing feature values against an individual’s unique baseline rather than using absolute population standards (Op De Beéck et al., 2018). This preprocessing step is essential to distinguish true condition-induced signals from a runner’s inherent movement style.

In conclusion, sleep deprivation represents a clinically relevant stressor for running populations, with established consequences for both performance and injury risk. Yet, despite the availability of both wearable technology and machine learning frameworks capable of detecting fatigue-related gait alterations, no study has examined whether sleep-deprivation-induced biomechanical changes during running are detectable using these tools. Therefore, the primary aim of this study was to determine whether sleep deprivation can be detected from running biomechanics using data from a single trunk-mounted inertial measurement unit (IMU). In addition, we will (i) compare the discriminative performance of a laboratory-based motion capture system (Vicon) and a wearable inertial sensor (Runeasi) in capturing sleep-deprivation-induced gait alterations, and (ii) evaluate the relative effectiveness of within-subject, paired classification and population-level classification approaches. Both modeling strategies were applied across motion capture systems, enabling a direct comparison between personalized models that leverage subject-specific baselines and generalized models that operate across individuals. We hypothesized that sleep deprivation would be detectable above chance level, with superior performance observed for within-subject classification compared to population-level models due to reduced inter-individual variability. Furthermore, although both motion capture systems were expected to demonstrate discriminative capacity, we anticipated that the laboratory-based motion capture system would outperform wearable inertial sensor owing to its richer and higher-resolution biomechanical feature space.

## 2 Materials and Methods

### 2.1 Participants

A total of 22 healthy, recreationally active runners (18–35 years) were initially recruited. Eligible participants were required to have adequate baseline sleep quality and duration, regular running habits (≥20 km/week for the past 6 months), and no running-related injuries in the preceding 2 months. Individuals with suspected sleep disorders, daily alcohol consumption, caffeine intake >400 mg/day, use of psychoactive substances, active smoking, or any known metabolic, cardiovascular, respiratory, or orthopedic conditions were excluded. One participant was subsequently excluded from analysis due to missing data. Descriptive characteristics are presented in Table 1. This study was approved by the Ethics Committee of UZ Leuven (S67790) and conducted in accordance with the Declaration of Helsinki. All participants provided written informed consent prior to participation.

**Table 1.** Subject characteristics.

|  | Male | Female |
| --- | --- | --- |
| Sample size | 10 | 11 |
| Age (years) | $22.1 \pm 2.2$ | $22.4 \pm 2.2$ |
| Body mass (kg) | $72.4 \pm 6.7$ | $59.6 \pm 8.2$ |
| Body height (m) | $1.80 \pm 0.06$ | $1.67 \pm 0.07$ |
| Km/week running | $36.7 \pm 25.2$ | $27.6 \pm 6.1$ |
| Speed at 1<sup>st</sup> lactate threshold (km/h) | $11.6 \pm 2.1$ | $10.1 \pm 1.2$ |
| Speed at 2<sup>nd</sup> lactate threshold (km/h) | $14.7 \pm 0.9$ | $12.6 \pm 0.8$ |
| Self-reported habitual sleep duration (h) | $8.0 \pm 0.9$ | $8.3 \pm 0.7$ |

### 3.2 Study Design

The full study design and experimental protocol are described in detail elsewhere (Seynaeve et al., 2026). Briefly, participants completed two experimental sessions in a randomized, counterbalanced crossover design: one following a night of normal sleep (control condition, CON) and one following a night of total sleep deprivation (SDEP), separated by a one-week washout period (Figure 1). In the CON condition, participants slept in the laboratory from 11:00 pm to 07:00 am under full polysomnographic monitoring. In the SDEP condition, participants remained awake throughout the night under continuous supervision. The running protocol commenced at 07:30 am following CON and at 06:30 am following SDEP, and consisted of three 8-minute intervals at 90% of LT1, LT1, and 90% of LT2, followed by a fourth interval at LT2 speed continued until volitional exhaustion. All running trials were performed on an instrumented treadmill (M-Gait, Motek, Amsterdam, the Netherlands). Running intensities were individually determined based on lactate thresholds established during a prior incremental treadmill test.

**Figure 1.**
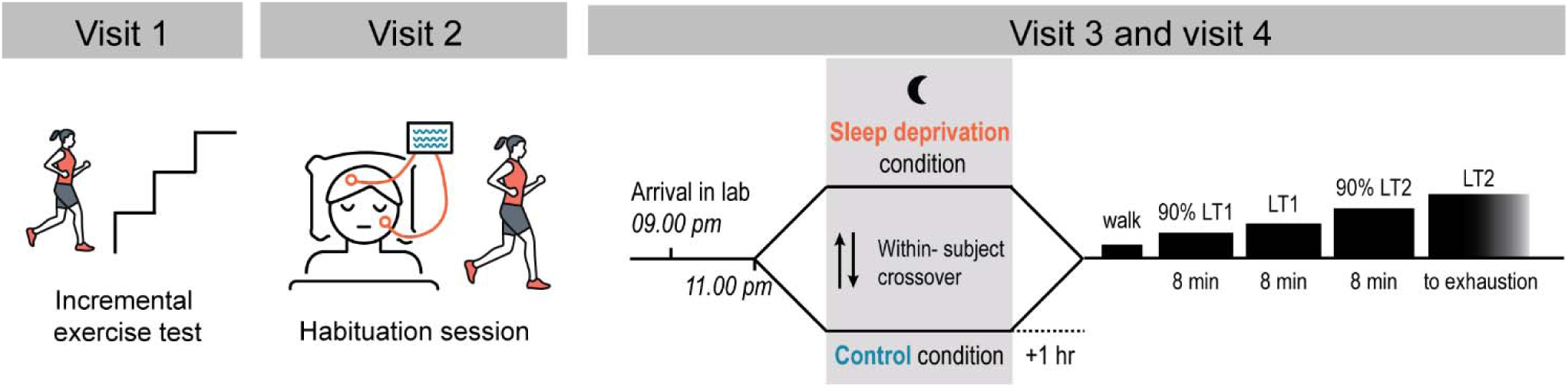
Schematic overview of the study visits. During visit 1, participants completed an incremental exercise test to determine individual lactate thresholds. Visit 2 served as a habituation session, familiarizing participants with the sleep laboratory and the running protocol. Visits 3 and 4 were completed in a counterbalanced, randomized order, consisting of one sleep deprivation night and one control night. The morning following each experimental night, participants performed three submaximal running intervals of 8 minutes at increasing intensities, followed by a single interval to volitional exhaustion at their second lactate threshold. LT1, first lactate threshold; LT2, second lactate threshold.

### 3.3 Instrumentation

Two motion capture systems were used simultaneously to record running biomechanics during each session. Laboratory-based kinematics were recorded using a 13-camera infrared motion capture system (Vicon, Oxford Metrics, Oxford, UK) sampling at 250 Hz, with 28 reflective markers. Concurrently, trunk accelerometry was recorded using the Runeasi system, a commercially available wearable sensor (Movesense, Vantaa, Finland) with integrated cloud-based processing platform (Runeasi.ai, Tienen, Belgium), sampling at 200 Hz. The sensor was mounted at the lower back between L5 and S2 using a firm elastic belt approximating the center of mass. Both systems recorded simultaneously throughout all running intervals, allowing direct comparison of their discriminative capacity under identical experimental conditions.

### 3.4 Feature Extraction

For the lab-based motion capture system, biomechanical features were extracted from the spatiotemporal, kinematic, kinetic and joint power variables described in detail elsewhere (Seynaeve et al., 2026). Variables were computed over strides collected between minutes 5 and 7 of the first three intervals and minutes 1 to 3 of the final interval. Only features normalized to body weight were used in the analysis. A total of 83 features were included in the final dataset.

For the wearable sensor, raw acceleration signals were automatically processed by the Runeasi platform into 21 running metrics computed in 5-second epochs. Epochs containing data corresponding to minutes 5 to 7 of the first three intervals and minutes 1 to 3 of the final interval were summarized by computing the mean, standard deviation, and left-right asymmetry of each metric, yielding 36 features per interval per participant. Left–right asymmetry was calculated as the absolute, non-directional percentage difference between limbs, ensuring that only the magnitude of inter-limb differences was retained. The extracted metrics included spatiotemporal characteristics (cadence, flight ratio, flight time, ground contact time), impact-related variables (impact magnitude, impact duration, peak loading rate, cumulative impact), trunk movement qualities (dynamic stability, smoothness), and anteroposterior/vertical ground reaction forces (braking, braking impulse, propulsive impulse, vertical impulse). A full description of each metric and its computation is provided in Supplementary Materials S1.

### 3.5 Classification Task

Two complementary classification tasks were evaluated. The first was a within-subject task, in which paired data from CON and SDEP sessions of a single participant were presented to the classifier. Then, the classifier was required to determine which session corresponded to each condition. This formulation explicitly removes inter-subject variability and isolates the intra-individual gait change due to sleep deprivation. The second was a between-subject task in which raw session measurements were classified as CON or SDEP without pairing, requiring the classifier to generalize across the natural variation in running gait between individuals. The between-subject task is more challenging and more generally applicable in a real-world setting, where a reference recording under rested conditions may not be available. Both approaches were applied independently to features derived from the lab system and from the wearable sensor, yielding four classification pipelines in total. In all cases, a single group-level model was trained across participants rather than subject-specific models.

### 3.6 Classification Models

Five supervised classification algorithms were evaluated: Logistic Regression, Decision Tree, Random Forest, k-Nearest Neighbours (k-NN), and Extreme Gradient Boosting (XGBoost). Each was subjected to systematic hyperparameter optimization using a nested cross-validation scheme, with the number of retained features treated as a tunable hyperparameter and optimized jointly with the model-specific parameters. Analyses were conducted separately for the two motion capture systems (lab and wearable) and for both classification tasks (within- and between-subject). All analyses were implemented in Python using *scikit-learn*, *XGBoost*, and *statsmodels*. A fixed random seed (42) was used throughout for reproducibility.

### 3.7 Data Preparation

#### Within-subject task

For each motion capture system, subject and running speed, biomechanical features from the CON and SDEP sessions were subtracted to produce two signed difference vectors: CON − SDEP and SDEP − CON. This way, each subject contributed two data points per speed condition. This formulation eliminates between-subject baseline differences and constrains the problem to detecting the directional shift in gait attributable to sleep deprivation.

#### Between-subject task

The same subjects and speed conditions were used, but the raw session measurements were retained without pairing. Each observation was labelled as CON or SDEP, and the classifier was required to generalize across subjects without access to a paired reference.

For both tasks, features were organized into a feature matrix X, a binary outcome vector y, and a group vector indicating subject identity. Running speed was encoded as a categorical variable with four levels (1-4), and used as a covariate in the ANOVA-based feature selection step.

### 3.8 Processing Pipeline

For each cross-validation fold, a three-stage pipeline was applied strictly within the training partition to prevent data leakage. *First*, all features were standardized to zero mean and unit variance using a standard scaler fitted on training data only. *Second*, an feature selector based on an analysis of variance (ANOVA) identified the features most strongly associated with condition type. Specifically, for each candidate feature, a two-way ANOVA (Type II) was fitted with condition and running speed as factors, including their interaction term. Features were ranked by the *p*-value of the condition main effect and the top *k* retained, where *k* was treated as a hyperparameter. *Third*, the selected classifier was trained on the standardized, reduced feature set.

### 3.9 Nested Cross-Validation and Hyperparameter Optimization

A nested cross-validation framework was used to provide an unbiased estimate of generalization performance. The outer loop used 10-fold group cross-validation (GroupKFold) stratified by subject identity, ensuring no subject appeared in both training and test partitions within a fold. For each outer training set, an inner GroupKFold grid search (up to five folds, capped by available subjects) exhaustively evaluated all hyperparameter combinations using accuracy as the criterion, and the best configuration was refitted on the full outer training set before evaluation on the held-out fold. Classification accuracy was recorded per outer fold and summarized as mean ± standard deviation across ten folds. A chance-level baseline of 50% applies to both tasks given the balanced binary design.

Hyperparameter search spaces were as follows. Logistic Regression: regularization strength *C* ∈ {0.001, 0.01, 0.1, 1, 10, 100}, *liblinear* solver. Decision Tree: maximum depth ∈ {2, 3, 4, unlimited}, minimum samples to split ∈ {2, 4, 6}. Random Forest: number of trees ∈ {50, 100, 200}, maximum depth ∈ {2, 3, unlimited}. k-NN: neighbors ∈ {3, 5, 7}, weighting ∈ {uniform, distance}. XGBoost: boosting rounds ∈ {5, 10, 50, 100}, maximum depth ∈ {2, 3, unlimited}, learning rate ∈ {0.01, 0.1, 0.2}. For all models, the number of ANOVA-selected features *k* was searched over {5, 10, 20, all}.

### 3.10 Feature Importance Analysis

To identify which gait features drove classification, each model was refitted on the full dataset and feature importances were extracted: absolute logistic regression coefficients for Logistic Regression, and Gini-based impurity reduction for Decision Tree, Random Forest, and XGBoost. k-NN does not yield intrinsic feature importances and was excluded from this analysis. Feature importances were computed separately for each sensor system and classification task.

## 3 Results

### 3.1 Within-subject classification

For the within-subject task, we classified the direction of the within-individual gait difference (CON − SDEP vs SDEP − CON). We found consistent above-chance performance across all models and both motion capture systems (Figure 2, Table 2). This confirms that sleep deprivation produces a systematic and directionally consistent change in running gait within individuals that is detectable by machine learning classifiers.

**Figure 2.**
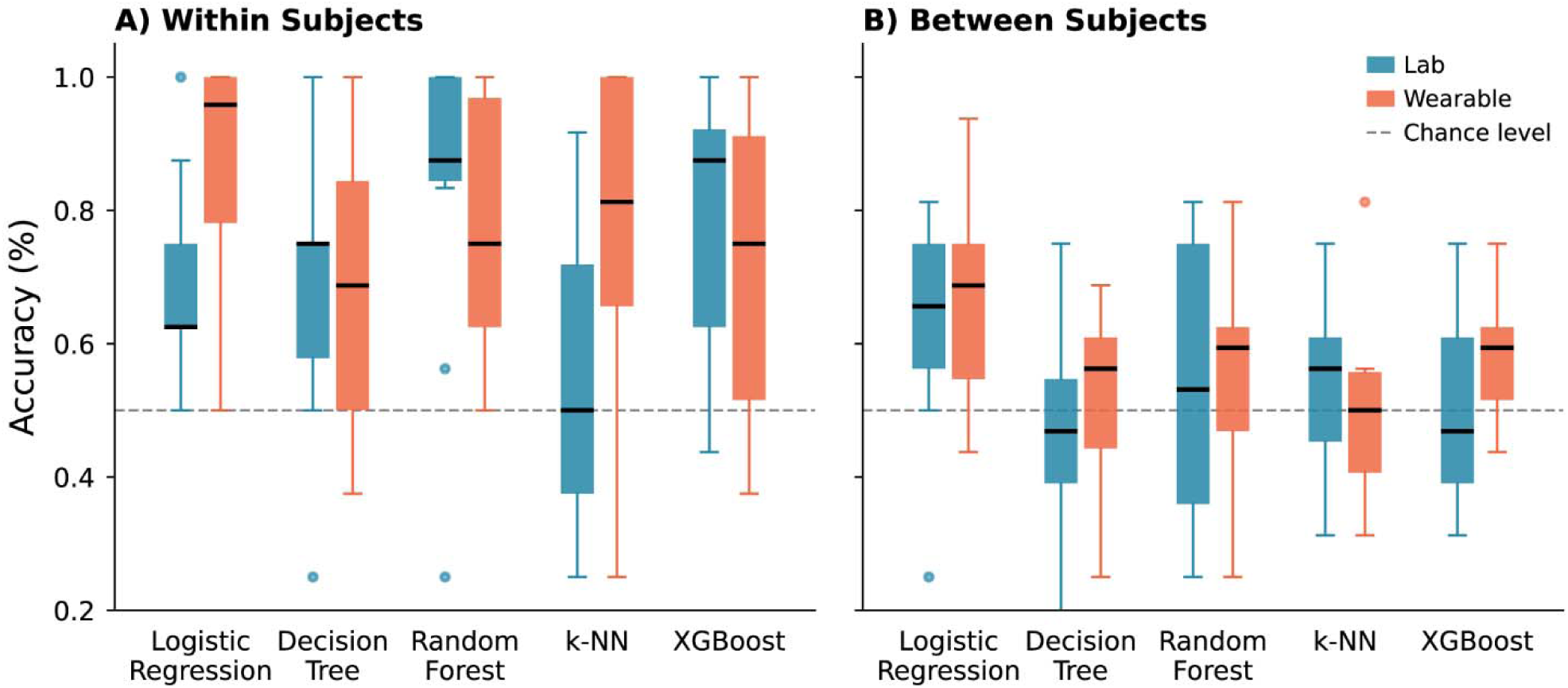
Classification accuracy of machine learning models for the within-subject (A) and between-subject task (B). Box plots showing cross-validated accuracy scores for five machine learning models evaluated on the lab-based (blue) and wearable (red) datasets. Each box represents the distribution of accuracy scores across cross-validation folds, with the horizontal line indicating the median, the box boundaries showing the interquartile range, and individual points representing outliers. The dashed horizontal line marks chance level (0.5). Both datasets were evaluated on identical model architectures with hyperparameter optimization performed via grid search. We found consistent above-chance performance across all models and both motion capture systems for the within-subject task but not for the between-subject task.

**Table 2.**
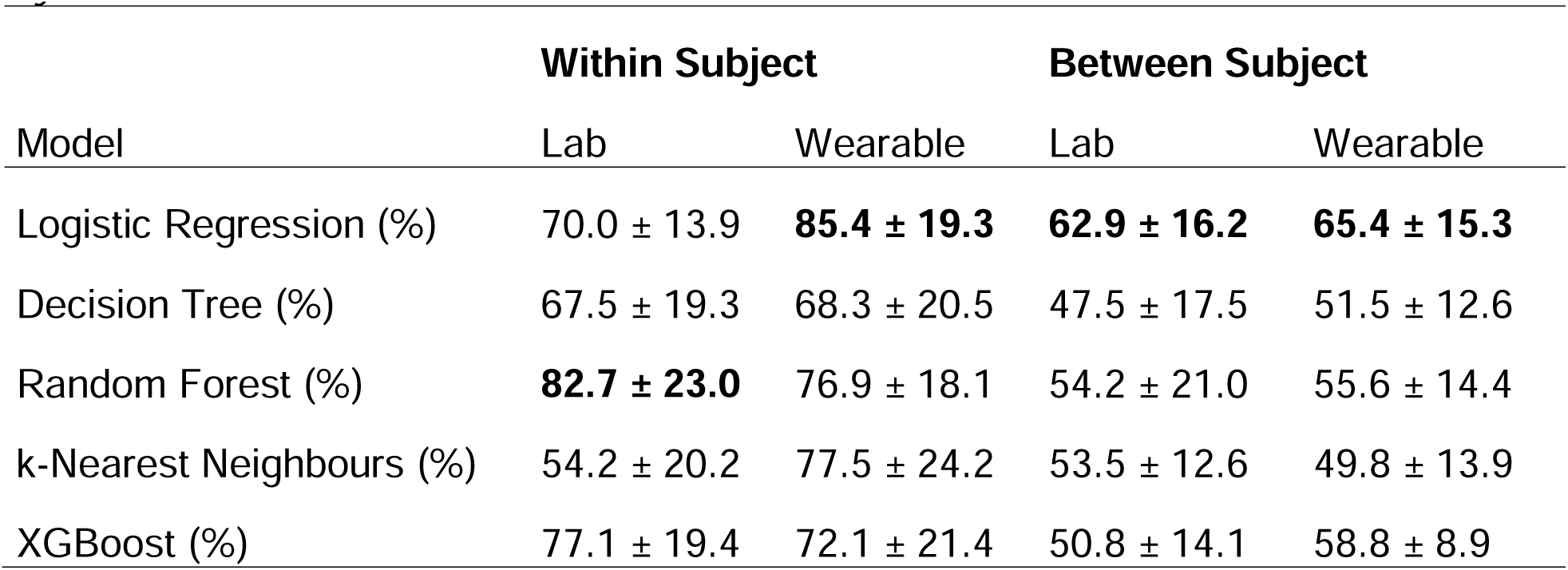
Classification accuracies for the different classification tasks and motion capture systems.

For the lab-based motion capture dataset, the ranking was markedly different. Random Forest led with a mean accuracy of 82.7% ± 23.0%, followed by XGBoost (77.1% ± 19.4%) and Logistic Regression (70.0% ± 13.9%). Decision Tree (67.5% ± 19.3%) performed at an intermediate level, while k-NN (54.2% ± 20.2%) fell considerably behind the other models. This pattern suggests that the richer, higher-dimensional motion capture feature space contains non-linear structure better exploited by ensemble tree methods, while distance-based approaches struggle in this higher-dimensional setting.

For the wearable sensor dataset, Logistic Regression achieved the highest mean accuracy (85.4% ± 19.3%), closely followed by k-NN (77.5% ± 24.2%) and Random Forest (76.9% ± 18.1%). Decision Tree (68.3% ± 20.5%) and XGBoost (72.1% ± 21.4%) performed at an intermediate level. The strong performance of the linear model again suggests that the within-subject gait difference captured by the wearable sensor is relatively low-dimensional and linearly separable, though the competitive performance of k-NN indicates a degree of local structure in the feature space as well.

Fold-level variability was substantial across both datasets. The best-performing model for the wearable dataset (Logistic Regression) showed a standard deviation of 19.3%, while the best-performing model for the lab-based dataset (Random Forest) showed a standard deviation of 23.0%, with some folds yielding perfect classification and others falling near or below chance. This indicates meaningful between-subject heterogeneity in the magnitude, though not the direction, of the gait response to sleep deprivation.

### 3.2 Between-subject classification

For the between-subject task, we predicted session type (CON or SDEP) from raw gait measurements without paired reference data. This task proved substantially more difficult, with all models performing close to or only modestly above the 50% chance level across both sensor systems (Figure 2, Table 2). This sharp decline relative to the within-subject task demonstrates that inter-subject variability in running gait dominates the sleep-deprivation signal when individuals are treated independently.

For the lab-based dataset, performance was constrained across all models. Logistic Regression performed marginally best (62.9% ± 16.2%), followed by k-NN (53.5% ± 12.6%), Random Forest (54.2% ± 21.0%), XGBoost (50.8% ± 14.1%), and Decision Tree (47.5% ± 17.5%).

For the wearable dataset, Logistic Regression again achieved the highest accuracy (65.4% ± 15.3%), representing the only model with a meaningful margin above chance. XGBoost (58.8% ± 8.9%) showed a modest but more consistent improvement over chance, notable for its comparatively low standard deviation. Random Forest (55.6% ± 14.4%) and Decision Tree (51.5% ± 12.6%) remained close to the chance baseline, while k-NN (49.8% ± 13.9%) performed at chance on average. Notably, Logistic Regression was the top performer on both motion capture systems for the between-subject task, in contrast to the within-subject task, where it ranked first for the wearable but third for the lab-based dataset. Nevertheless, no model achieved classification accuracy that would be interpretable as reliable detection of sleep deprivation at the group level without individual reference data.

### 3.3 Feature importance

Feature importances were computed on the full dataset for the within-subject task, providing insight into which gait characteristics are most sensitive to sleep deprivation at the individual level (Figure 3).

**Figure 3.**
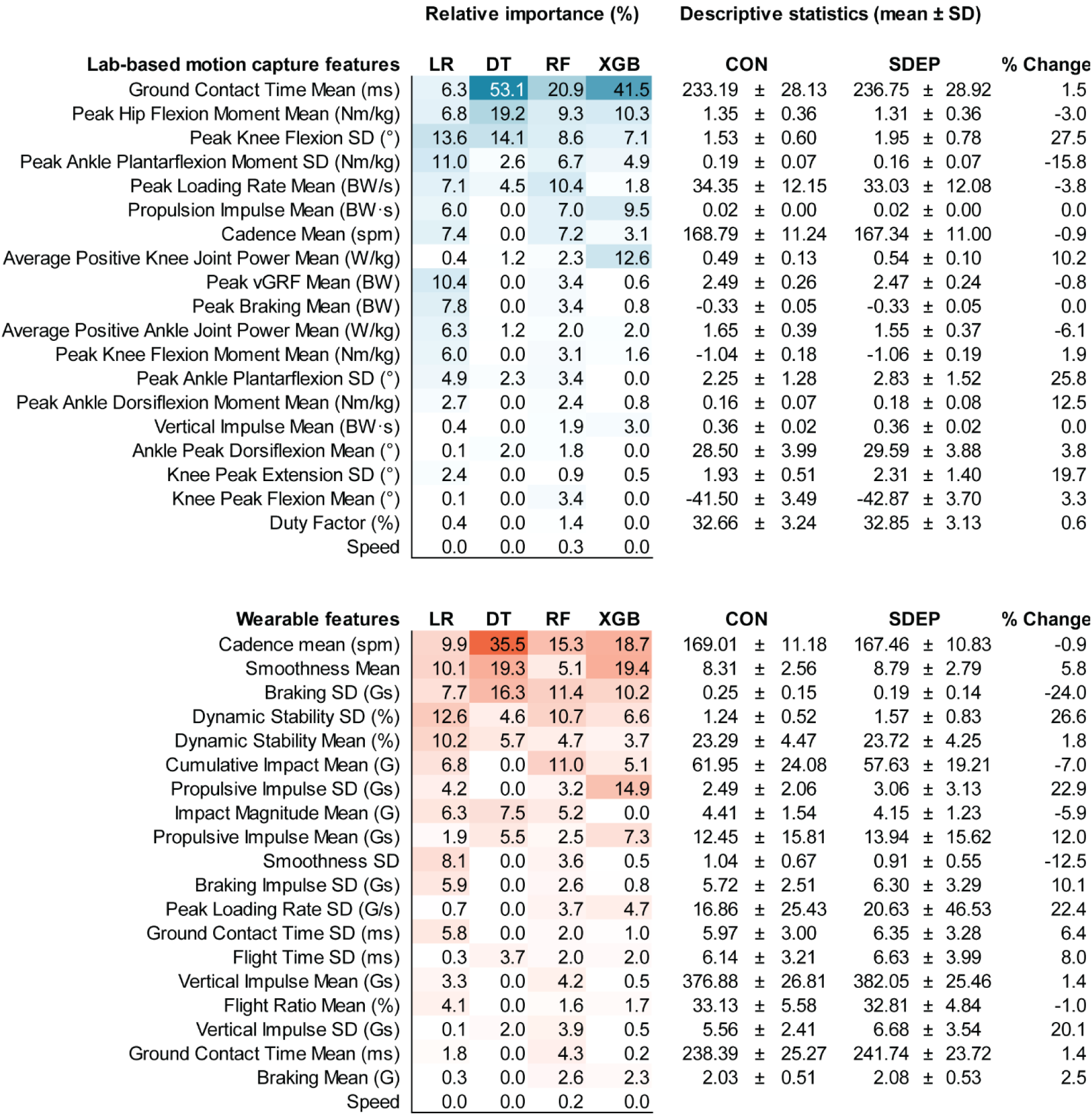
Feature importance heatmap across four classifier models for the within-subject task. Normalized feature importance (0–1) of (A) lab-based motion capture features (blue), and (B) wearable-derived features (red). Features are ranked by mean importance; darker shading = higher importance. Normalization was applied within each model. Across both systems, variability-based features (SD metrics) were consistently among the most important discriminators, alongside spatiotemporal features such as ground contact time and cadence. Importance was highly concentrated in a single feature for the decision tree, whereas Random Forest and XGBoost showed more distributed profiles across both the lab-based and wearable systems. LR, Logistic Regression; DT, Decision Tree; RF, Random Forest; XGB, XGBoost.

#### Lab-based dataset

Ground contact time was the dominant feature for tree-based models, ranking first for Decision Tree (53.1%), Random Forest (20.9%), and XGBoost (41.5%), though it contributed less distinctively in Logistic Regression, where importance was more evenly distributed across features. Peak hip flexion moment was the second most important feature overall, contributing consistently across all four models (6.8%, 19.2%, 9.3%, 10.3%). Peak knee flexion variability also appeared among the top features across all models (13.6%, 14.1%, 8.6%, 7.1%), followed by peak ankle plantarflexion moment variability and peak loading rate as moderately important contributors. Overall, the pattern points to sleep deprivation altering ground contact mechanics and lower-limb loading characteristics in a consistent within-individual direction, with kinetic and spatiotemporal variables jointly driving classification across the full-body motion capture feature space.

#### Wearable dataset

Cadence was the single most consistently important feature across models, ranking first for Logistic Regression (9.9%), Decision Tree (35.5%), and Random Forest (15.3%), and second for XGBoost (18.7%). Smoothness mean was similarly prominent, ranking first for XGBoost (19.4%) and second for Decision Tree (19.3%), and contributing meaningfully in Logistic Regression (10.1%). Braking SD emerged as a consistently important contributor across all four models (7.7%, 16.3%, 11.4%, 10.2%), as did Dynamic Stability SD (12.6%, 4.6%, 10.7%, 6.6%). Beyond these leading features, cumulative impact mean and propulsive impulse variability appeared as moderately important for Logistic Regression, Random Forest and XGBoost. The prominence of variability features alongside mean-level variables suggests that sleep deprivation affects not only the central tendency of key gait parameters but also the consistency of movement mechanics within individuals.

## 4 Discussion

This study examined whether total sleep deprivation produces detectable changes in running gait biomechanics. Analyses were performed using two complementary motion capture systems - a laboratory-based full-body system (Vicon) and a field-deployable trunk-mounted wearable (Runeasi) - and two classification tasks that differed depending on the availability of an individual reference condition. These analyses resulted in three primary findings. *First*, within-subject classification was consistently above chance for both motion capture systems, confirming that sleep deprivation produces a systematic and individually consistent biomechanical signature in running gait. *Second*, between-subject classification failed across nearly all models and both motion capture systems, demonstrating that inter-individual variability in running gait overwhelms the sleep-deprivation signal when no reference condition is available. *Third*, despite their fundamental differences in measurement scope, both systems converged on temporal, loading-related, and variability-based features as the most discriminative signals of sleep deprivation. These findings partially support our hypotheses: sleep deprivation was detectable above chance for the within-subject task, and within-subject classification outperformed between-subject classification as anticipated. However, contrary to our expectation that the lab-based motion capture would outperform the wearable by virtue of its richer feature space, both systems achieved comparable within-subject classification accuracy, and neither system supported meaningful between-subject detection.

### 4.1 Within-subject vs between-subject classification

Within-subject classification accuracy exceeded chance level across all models and both measurement modalities. This provides evidence that total sleep deprivation alters running gait in a sufficiently systematic and repeatable manner within individuals to support machine learning classification. These findings extend recent laboratory-based work demonstrating meaningful biomechanical changes following a single night of sleep deprivation by showing that such alterations are sufficiently structured to permit condition discrimination at the individual level (Seynaeve et al., 2026). However, fold-level variability was substantial across both systems, with individual folds ranging from near-perfect classification to near-chance performance. This likely reflects heterogeneity in the magnitude of the gait response to sleep deprivation across individuals, although variability in model estimation and fold composition may also have contributed. Such variability is nonetheless consistent with well-documented inter-individual differences in vulnerability to sleep deprivation (Van Dongen et al., 2004)

By contrast, between-subject classification failed to produce meaningful discrimination in either modality, with most models performing within a few percentage points of the 50% chance baseline. As documented extensively in the exercise-induced fatigue detection literature, inter-individual differences in gait patterns often exceed the magnitude of condition-induced effects (Chalitsios et al., 2024). The present findings suggest that this challenge extends to the sleep deprivation context. More broadly, these results support the subject-specific, baseline-normalized approach advocated in the fatigue literature (Op De Beéck et al., 2018), indicating that the absence of an individual reference measurement substantially limits the reliable detection of sleep-deprivation-related gait changes. In practical terms, a single gait recording obtained without knowledge of an individual’s rested running pattern appears insufficient to reliably infer sleep deprivation status.

### 4.2 Wearable sensors vs 3D motion capture

Contrary to our hypothesis that 3D motion capture would outperform a wearable system, both systems achieved comparable within-subject classification accuracy, and neither supported meaningful between-subject detection. However, the optimal model family differed notably between the two systems. Using wearable sensor features, linear and distance-based models (Logistic Regression and k-NN) outperformed tree-based ensemble methods (Random Forest and XGBoost), whereas the reverse pattern was observed for the lab-based motion capture features.

The interpretation of this dissociation warrants caution, as model performance alone cannot establish the underlying geometry of a feature space, and multiple explanations may contribute. One plausible contributor is the difference in feature dimensionality between the two candidate pools. Large-scale benchmarking across 243 datasets has shown that the number of features is significantly correlated with the performance advantage of Random Forest over Logistic Regression, with the advantage shifting from negative to positive as dimensionality increases, while sample size alone was not a significant predictor of this gap (Couronné et al., 2018). The lab-based motion capture system drew from a substantially larger and more heterogeneous candidate pool of 83 features spanning multiple biomechanical domains, compared to 36 features for the wearable system. If the effective dimensionality and structural complexity of the retained feature sets differed accordingly, this could partly explain why ensemble methods outperformed linear models for the lab-based system but not for the wearable system.

A further, non-mutually-exclusive explanation concerns feature intercorrelation introduced by the selection procedure. Because ANOVA-based univariate feature selection ranks features by their individual association with the outcome without accounting for redundancy among predictors (Michalak and Kwasnicka, 2010; Walowe Mwadulo, 2016), features selected from the larger, more heterogeneous lab-based pool may carry greater residual intercorrelation. Whether this differentially affected the two model families is difficult to establish without formal analysis of inter-feature correlations in the retained subsets, and this explanation should be treated as a plausible hypothesis rather than a definitive account of the observed performance differences.

### 4.3 Feature importance

Feature importances were computed on the full dataset for the within-subject task to identify which features contributed most strongly to discrimination between rested and sleep-deprived conditions at the individual level. Despite substantial differences in feature representation between the two systems, comparison of importance patterns revealed convergence across three recurring themes. The most pronounced concerned temporal structure: cadence was the dominant wearable feature across models, while ground contact time emerged as the dominant laboratory-based feature. These variables represent closely related components of stride-cycle timing. Therefore, their co-emergence as highly discriminative features across independent motion capture systems suggests that alterations in the temporal organization of running were among the most consistently discriminative signals of sleep deprivation.

A second recurring theme involved loading-related information. Impact-related wearable features, including cumulative impact and impact magnitude, and laboratory-derived kinetic variables, including peak loading rate and peak braking force, all contributed meaningfully to classification. While trunk accelerometry provides an indirect representation of impact transmission, laboratory-based kinetics directly characterize ground reaction force mechanics. Therefore, these variables reflect related but not equivalent aspects of impact behavior. Nevertheless, their parallel importance suggests that loading-related information contributed substantially to discrimination across measurement contexts.

A third shared pattern was the consistent contribution of variability-related features alongside mean-level variables in both systems. This indicates that stride-to-stride variability in key gait characteristics contributed meaningfully to discrimination between rested and sleep-deprived running states. Importantly, however, importance rankings alone cannot establish the direction of these changes; variability-related features may contribute strongly to classification regardless of whether variability increased, decreased, or shifted in a more complex multivariate manner.

The two systems diverged in ways that reflect their different measurement scope. Wearable importance was distributed more diffusely across trunk-derived variables including dynamic stability, smoothness, and multiple variability metrics, whereas laboratory-based importance concentrated on specific joint-level kinetic and kinematic variables, reflecting the system’s capacity to resolve mechanical detail inaccessible to a single trunk-mounted sensor. This suggests the two modalities may provide different but partially overlapping perspectives on the gait response to sleep deprivation, with the wearable capturing global coordination and trunk movement organization and the laboratory system isolating joint-specific adaptations.

The patterns described above are consistent with several hypothesized neuromuscular consequences of sleep deprivation, and find support in the descriptive data. First, impaired feedforward control may play a central role. Leg stiffness at ground contact depends in part on anticipatory muscle preactivation, governed by feedforward mechanisms that allow the nervous system to predict upcoming mechanical demands and adjust muscle activity accordingly (Müller et al., 2010, 2014; Khajooei et al., 2024). Sleep deprivation impairs higher-order sensorimotor and predictive control processes (Killgore, 2010; Umemura et al., 2021), which may reduce the precision with which muscle activation is timed and scaled before ground contact. This may leave the leg inadequately stiffened at touchdown, potentially contributing to the longer ground contact times, reduced cadence, and lower impact-related variables observed across both systems. Second, the shift toward reduced loading may reflect a conservative protective strategy in which load reduction is prioritized over stiffness and propulsive efficiency when precise anticipatory control is compromised, a pattern previously proposed in the context of extreme endurance running (Vernillo et al., 2014; Degache et al., 2016). Third, the prominence of hip and knee kinetics alongside reduced ankle joint power in the lab-based dataset suggests a possible distal-to-proximal redistribution of joint work. During running, plantarflexors operate at a disproportionately high proportion of their maximal force and contribute a larger share of positive work than other muscle groups (Dorn et al., 2012; Kulmala et al., 2016), making them particularly susceptible to fatigue. A compensatory distal-to-proximal shift has been observed during prolonged running to exhaustion (Sanno et al., 2018; Willer et al., 2021), and it is plausible that centrally mediated fatigue under sleep loss may produce a similar pattern, though this remains speculative in the absence of direct electromyographic evidence. Finally, stride-to-stride inconsistency seemingly increased across the majority of kinematic and wearable-derived variability metrics. This is consistent with evidence that sleep deprivation impairs the regularity of continuous coordinated movement, likely through degraded sensorimotor feedback and executive control processes (Umemura et al., 2021, 2022; Paillard, 2023).

However, it should be acknowledged that feature importance values reflect contribution to model discrimination rather than direct physiological sensitivity, and the following interpretations should be read as hypothesis-generating rather than conclusive. These mechanisms are not mutually exclusive, and because importance measures are operationalized differently across algorithms and are sensitive to feature covariance and model architecture, the patterns reported here should be interpreted as suggestive of recurring discriminative themes rather than definitive mechanistic conclusions.

### 4.4 Limitations

Several limitations of the present study should be acknowledged. The sample size of 21 participants, while typical for laboratory-based gait studies and adequate for the within-subject design, limits statistical power and may contribute to the fold-level variability observed in cross-validation (Vabalas et al., 2019). Replication in larger samples is required to establish the robustness and generalizability of the findings.

The stability of selected features and model performance across resampling folds or participant subsets was not explicitly quantified, and the feature importance patterns reported should therefore be interpreted with caution. Feature importances were computed by refitting each model on the full dataset rather than aggregating importance estimates across held-out folds. This provides a descriptive overview of which features contribute most to in-sample classification, but the resulting estimates are exploratory in nature and not derived from independent out-of-sample evaluation. Moreover, importance measures are inherently model-dependent, differing across algorithms through coefficients, impurity-based measures, or gain metrics, and are sensitive to feature covariance, redundancy, and model structure (Nicodemus et al., 2010; Toloşi and Lengauer, 2011). As such, they should not be interpreted as direct indicators of physiological sensitivity.

The experimental protocol involved total sleep deprivation, which represents an extreme and relatively uncommon condition in real-world athletic settings. More ecologically prevalent forms of sleep restriction, such as partial sleep loss over multiple nights, may produce subtler or qualitatively different gait adaptations. It therefore remains uncertain to what extent the present findings generalize to more chronic or realistic sleep loss scenarios.

The wearable system (Runeasi) processes raw accelerometry through a proprietary algorithm, and the exact computational definitions of several derived metrics are not fully transparent. This limits mechanistic interpretability and reduces cross-platform reproducibility of specific features.

Finally, all running trials were performed on a treadmill at fixed speeds. Although this constrains ecological validity, it was a deliberate methodological choice: treadmill running allowed for precise speed control, which is essential given that speed directly influences cadence and other spatiotemporal parameters. It therefore remains unclear whether the observed gait changes would manifest similarly during overground running or at self-selected speeds, where pacing strategies may shift in response to fatigue and speed can no longer be held constant.

Taken together, these limitations highlight the need for cautious interpretation of the present findings and for future work to evaluate their robustness across larger samples, more ecologically valid sleep loss conditions, and more naturalistic running environments.

## 5 Conclusions

This study demonstrates that total sleep deprivation induces a detectable and individually consistent biomechanical signature in running gait, identifiable by both a laboratory motion capture system and a field-deployable trunk-mounted wearable. Within-subject classification performed consistently above chance across both systems, whereas between-subject classification failed, confirming that individual baseline data are essential for reliable gait-based fatigue detection. This conclusion mirrors findings from the exercise-induced fatigue literature and extends them to a sleep deprivation context. Contrary to our hypothesis, the lab-based motion capture system did not outperform the wearable device. Instead, both systems captured complementary rather than redundant aspects of the gait response. Together, these findings provide a biomechanical and methodological foundation for wearable-based fatigue monitoring in runners, and highlight longitudinal, individualized monitoring as a necessary approach for detecting sleep-deprivation effects in real-world running contexts.

## 6 Conflict of Interest

Benedicte Vanwanseele is a co-founder and shareholder of Runeasi, the manufacturer of the wearable sensor used in this study. Kilian Hendrickx is currently employed by Runeasi but was not employed by the company during the design of the study, data collection, data analysis, or interpretation of the results. All analyses were completed prior to the start of his employment. The remaining authors declare no commercial or financial relationships that could be construed as a potential conflict of interest.

## 7 Author Contributions

**Maura Seynaeve:** Conceptualization; Methodology; Investigation; Data curation; Formal analysis; Visualization; Project administration; Writing – original draft; Writing – review & editing.

**Kilian Hendrickx:** Software; Formal analysis; Methodology.

**Benedicte Vanwanseele**: Writing – review & editing; Supervision.

**Toon De Beukelaar:** Writing – review & editing; Supervision; Funding acquisition.

## 8 Funding

This work was supported by the KU Leuven Special Fund (grant no. 3M220016 to T.d.B.).

## 9 Acknowledgments

We would like to thank runeasi.ai for providing the materials and platform used to collect and process the accelerometer data.

## 10 Data Availability Statement

The datasets generated and/or analyzed in this study are not publicly available due to ethical considerations and participant privacy restrictions. Requests to access the data should be directed to the corresponding author and the Social and Societal Ethics Committee at KU Leuven.

