## Supplementary material for "Detecting Sleep Deprivation from Running Biomechanics Using Machine Learning Classification: A Comparison Between Wearable and Laboratory Motion Capture": Table_S1

| **Feature** | **Mean** | **SD** | **Asymmetry** | **Description** | **Unit** |
| --- | --- | --- | --- | --- | --- |
| ***Spatiotemporal*** | | | | | |
| Cadence | ✕ |  |  | Number of steps per minute. | steps/min |
| Flight ratio | ✕ |  |  | Ratio between the flight time and total step duration. | % |
| Flight time | ✕ | ✕ | ✕ | Total duration the foot is in the air during the swing phase. Asymmetry expressed as percentage difference between left and right leg. | ms / % |
| Ground contact time | ✕ | ✕ | ✕ | Total duration the foot is in contact with the ground. Asymmetry expressed as percentage difference between left and right leg. | ms / % |
| ***Impact*** | | | | | |
| Impact magnitude | ✕ | ✕ | ✕ | Vertical acceleration peak that reaches the pelvis immediately after ground contact, averaged across left and right leg. Asymmetry expressed as percentage difference between left and right leg. | G / % |
| Impact duration | ✕ | ✕ | ✕ | Time for the impact shockwave to travel from the foot to the pelvis; reflects shock absorption capacity of the legs. Longer durations indicate better absorption. Asymmetry expressed as percentage difference between left and right leg. | ms / % |
| Peak loading rate | ✕ | ✕ | ✕ | Rate of change of the vertical impact acceleration at initial ground contact, averaged across left and right leg. Asymmetry expressed as percentage difference between left and right leg. | G/s / % |
| Cumulative impact | ✕ |  |  | Accumulated vertical impact load over the recording interval. | G |
| ***Ground reaction forces*** | | | | | |
| Braking | ✕ | ✕ | ✕ | Anteroposterior braking acceleration that reaches the pelvis immediately after foot strike, averaged across left and right leg. Asymmetry expressed as percentage difference between left and right leg. | G / % |
| Braking impulse | ✕ | ✕ | ✕ | Sum of all negative anteroposterior acceleration values during the stance phase, averaged across left and right leg. Asymmetry expressed as percentage difference between left and right leg. | Gs / % |
| Propulsive impulse | ✕ | ✕ | ✕ | Sum of all positive anteroposterior acceleration values during the stance phase, averaged across left and right leg. Asymmetry expressed as percentage difference between left and right leg. | Gs / % |
| Vertical impulse | ✕ | ✕ | ✕ | Sum of all vertical acceleration values during the stance phase, averaged across left and right leg. Asymmetry expressed as percentage difference between left and right leg. | Gs / % |
| ***Trunk movement quality*** | | | | | |
| Dynamic stability | ✕ | ✕ | ✕ | Proportion of hip movement in the medio-lateral direction relative to overall movement during landing, averaged across left and right leg. Higher values indicate more lateral energy loss. Asymmetry expressed as percentage difference between left and right leg. | % / % |
| Smoothness | ✕ | ✕ | ✕ | Jerk cost calculated as the area under the anteroposterior jerk curve during the stance phase, averaged across left and right leg. Lower values indicate smoother force application. Asymmetry expressed as percentage difference between left and right leg. | a.u. / % |
